# Progranulin maintains blood pressure and vascular tone dependent on EphrinA2 and Sortilin1 receptors and eNOS activation

**DOI:** 10.1101/2023.04.02.534563

**Authors:** Ariane Bruder-Nascimento, Wanessa M.C. Awata, Juliano V. Alves, Shubhnita Singh, Rafael M. Costa, Thiago Bruder-Nascimento

**Affiliations:** Department of Pediatrics; Center for Pediatrics Research in Obesity and Metabolism (CPROM); Endocrinology Division at UPMC Children’s Hospital of Pittsburgh; Vascular Medicine Institute (VMI), University of Pittsburgh, Pittsburgh, PA, USA

**Keywords:** progranulin, blood pressure, vascular function, nitric oxide

## Abstract

**Background:** The mechanisms determining vascular tone are still not completely understood, even though it is a significant factor in blood pressure management. Many circulating proteins have a significant impact on controlling vascular tone. Progranulin (PGRN) displays anti-inflammatory effects and has been extensively studied in neurodegenerative illnesses. We investigated whether PGRN sustains the vascular tone that helps regulate blood pressure.

**Methods:** We used male and female C57BL6/J wild type (PGRN+/+) and B6(Cg)-Grn^tm1.1Aidi^/J (PGRN-/-) to understand the impact of PGRN on vascular contractility and blood pressure.

**Results:** We found that male and female PGRN-/- mice display elevated blood pressure followed by hypercontractility to noradrenaline in mesenteric arteries, which are restored by supplementing the mice with recombinant PGRN (rPGRN). In *ex vivo* experiments, rPGRN attenuated the vascular contractility to noradrenaline in male and female PGRN+/+ arteries, which was blunted by blocking EphrinA2 or Sortlin1. To understand the mechanisms whereby PGRN evokes anti-contractile effects, we inhibited endothelial factors. L-NAME [nitric oxide (NO) synthase (NOS) inhibitor] prevented the PGRN effects, whereas indomethacin (cyclooxygenases inhibitor) only affected the contractility in arteries incubated with vehicle, indicating the PGRN increases nitric oxide and decreases contractile prostanoids. Finally, rPGRN induced endothelial NOS (eNOS) phosphorylation and NO production in isolated mesenteric endothelial cells.

**Conclusion:** Circulating PGRN regulates vascular tone and blood pressure via EphrinA2 and Sortlin1 receptors and eNOS activation. Collectively, our data suggest that deficiency in PGRN is a cardiovascular risk factor and that PGRN might be a new therapeutic avenue to treat high blood pressure.

*Clinical Perspective:* What is new? - PGRN displays vascular anti-contractile effects dependent on EphrinA2 and Sortilin1 receptors and nitric oxide formation in male and female
- Deficiency in PGRN triggers high blood pressure and induces vascular dysfunction characterized by hypercontractility to noradrenaline
- PGRN supplementation restores blood pressure and vascular dysfunction in PGRN-deficient mice What are the clinical implications? - PGRN deficiency is associated with neurodegenerative diseases including neuronal ceroid lipofuscinosis and frontotemporal dementia (FTD). Our study reveals that a lack of PGRN might be associated with vascular dysfunction and high blood pressure
- Supplementation with PGRN might be a potential therapeutic route to treat high blood pressure and diseases associated with vascular dysfunction
- Reduction in PGRN might be a target to screen for higher cardiovascular risk

## Introduction

Hypertension, commonly known as high blood pressure (HBP), is a chronic medical condition that affects millions of people worldwide. It is a silent killer that often goes undetected for years, and if left untreated, can lead to serious health consequences such as heart disease, stroke, and kidney failure. HBP is still regarded as the primary risk factor for the global burden of illness despite substantial advancements in prevention and treatment^1, 2^, and it equally affects people in economically developed and developing countries^1, 3^. HBP patients are more likely to suffer from chronic renal disease, dementia, myocardial infarction, and stroke^2, 3^. The central nervous system (CNS), the kidneys, and the vasculature have all been identified as potential triggers for HBP^3–7^. In this work, our main goal was to comprehend how changes in the vasculature contribute to the development of HBP.

Vascular resistance is supported by three major factors: the viscosity of the blood, the length, and the diameter of the blood vessels^1^. High systemic vascular resistance, which can lead to HBP, is primarily caused by changes in vascular stiffness, inflammation, and tone^3, 5, 6, 9–11^. Control of vascular tone is coordinated primarily by a balance between endothelium-derived relaxing factors (EDRF) such as nitric oxide, prostaglandins, endothelium-derived hyperpolarizing factor (EDHF), and endothelium-derived contracting factors (EDCF) such as endothelin-1, thromboxane, and reactive oxygen species^2–5^, an unbalance between these factors triggers vascular hypercontractility leading to elevated vascular resistance.

Obesity^6, 7^, diabetes^8, 9^, and lipodystrophy^10–12^ are major causes of HBP. Interestingly, these conditions share a striking circulating increase of progranulin (PGRN)^13–17^, however why PGRN is elevated in the plasma and if this increase can modulate the vascular tone and blood pressure is unknown. PGRN is a highly conserved glycoprotein that is expressed and secreted by adipocytes, neurons, immune cells, and endothelial cells^18–20^. It plays a pivotal role in regulating wound healing, cell growth, and inflammation via autocrine, paracrine, or endocrine actions^18–20^. Studies have shown that PGRN has anti-inflammatory and antihypertrophic aspects in sepsis^21^, liver fibrosis^22^, diabetic nephropathy^15, 23^, age-related cardiac disorders^24^, ischemia/reperfusion diseases^25^, and atherosclerosis^26–28^. Other studies have found that PGRN is associated with neurodegenerative diseases^18, 29, 30^. For instance, loss of PGRN causes neuronal ceroid lipofuscinosis ^19, 31^ and frontotemporal dementia (FTD)^19, 31^, while reduced PGRN levels increase the risk of Parkinson’s disease and Alzheimer’s disease^19^. Finally, prior studies have shown a connection between PGRN and endothelial activity via regulating protein kinase B (AKT) and endothelial nitric oxide synthase (eNOS), as well as blocking nuclear factor-κB (NFkB)^26, 32, 33^. Although a connection between endothelial activation and PGRN has been previously suggested, its association with vascular tone and blood pressure is still undetermined.

Herein, we used PGRN-deficient mice and recombinant PGRN (rPGRN) combined with pharmacological interventions to better understand the contribution of circulating PGRN to vascular tone and blood pressure regulation and more specifically to test the hypothesis that PGRN maintains vascular function and blood pressure via modulating endothelial factors.

## Material and methods

The data that support the findings of this study are available from the corresponding author upon reasonable request.

Eleven-to thirteen-week-old male and female C57BL6/J wild type (PGRN+/+) and B6(Cg)- Grn^tm1.1Aidi^/J (PGRN-/-) were used. All mice were fed with standard mouse chow and tap water was provided ad libitum. Mice were housed in an American Association of Laboratory Animal Care–approved animal care facility in the Rangos Research Building at the Children’s Hospital of Pittsburgh of the University of Pittsburgh. Institutional Animal Care and Use Committee approved all protocols (IACUC protocols # 19065333 and 22061179). All experiments were performed in accordance with Guide Laboratory Animals for The Care and Use of Laboratory Animals.

### Mouse models of HBP

To characterize the circulating levels of PGRN in HBP, we used two different models.

- *Angiotensin II (Ang-II) treated mice^34^*: Male mice were infused with vehicle or Ang-II (490 ng/min/kg) for 14 days with ALZET osmotic minipumps (Alzet Model 1002; Alzet Corp Durect, Cupertino, CA) while receiving regular drinking water.
- *Aldosterone (Aldo) treated mice^35^*: Male mice were infused with vehicle or aldosterone (600 μg/kg per day) for 14 days with ALZET osmotic minipumps (Alzet Model 1002; Alzet Corp Durect, Cupertino, CA) while receiving 1% saline in the drinking water.

### Sodium urine collection and measurement

Changes in blood pressure affect the urine volume and sodium excretion, therefore we placed PGRN+/+ and PGRN-/- mice placed in a metabolic cage for 24h-urine collection. Briefly, mice were acclimated for 24h followed by an additional 24h for sample collection and sodium measurement. Urine was sent to Kansas State Veterinary Diagnostic Laboratory for sodium quantification.

### Restoration of circulating PGRN

Mice were treated with rPGRN ALZET osmotic minipumps (Alzet Model 1001; Alzet Corp Durect, Cupertino, CA) for 7 consecutive days (20ug/day), as described elsewhere^13^.

### Circulating PGRN levels

Enzyme-Linked Immunosorbent Assay (ELISA) was used to detect plasmatic PGRN (R&D biosystem).

### Circulating inflammatory profile via proteome profiler mouse cytokine

Plasma from PGRN+/+ and PGRN-/- mice was collected and frozen at − 80 C until the day of the analysis. Plasma from six mice of each group was pooled and the experiment was performed in duplicate. The proteome profiler mouse cytokine array kit (R&D System) was performed following the manufacturer’s instructions. The intensity of each band generated in the assay was analyzed in ImageJ. Data were presented in fold changes in a Heat Map graph.

### *In vivo* blood pressure measurement

Blood pressure was analyzed via radiotelemetry using an HD-X10 telemeter (Data Sciences International, St Paul, MN). Transmitters were implanted as described previously^36, 37^. After 7 days of recovery from surgery, necessary for the mice to gain their initial body weight, data were recorded for 5 days as a baseline. Then, rPGRN was continuously administrated as described above. Systolic blood pressure (SBP), diastolic Blood Pressure (DBP), mean arterial pressure (MAP), and heart rate were analyzed.

### Indices of autonomic function

To analyze whether changes in autonomic response interfere with any changes in blood pressure, indices of the autonomic function were obtained on the last day of the recording baseline period. A classic pharmacological method consisting of a single intraperitoneal injection of the ganglionic blocker mecamylamine (5 mg/kg) or of the β-adrenergic receptor blocker propranolol (6 mg/kg) was used^36–38^. Injections were conducted more than 2 h apart in random order. Changes in blood pressure response to pharmacological compounds within 60 min post-injection were reported. Data were expressed as a percentage of the baseline value.

### Histology

PGRN+/+ and PGRN-/- were euthanized for aortae harvest and perfused with cold phosphate-buffered saline (PBS). Aortae were collected and placed in a 4% paraformaldehyde (PFA) solution for histology analysis. After 12h in PFA, tissues were placed in 70% ethanol until the day of preparing the samples for histology. Aortae were embedded in paraffin, then samples were sectioned and stained with hematoxylin and eosin (H&E) and Masson’s Trichrome to analyze the vascular remodeling and structure.

### Vascular function

Rings from second-order mesenteric resistance arteries were mounted in a wire myograph (Danysh MyoTechnology) for isometric tension recordings with PowerLab software (AD Instruments) as described before^34, 35, 39^. Briefly, rings (2mm) were placed in tissue baths containing warmed (37 °C), aerated (95% O_2_, 5% CO_2_) Krebs Henseleit Solution: (in mM: 130 NaCl, 4.7 KCl, 1.17 MgSO_4_, 0.03 EDTA, 1.6 CaCl2, 14.9 NaHCO_3_, 1.18 KH_2_PO_4_, and 5.5 glucose) and after 30 min of stabilization, curves of tension were performed to adjust the ideal tension for each segment followed by incubation with KCl (60mM).

- *Protocol to study the effects of PGRN:* Since noradrenaline is endogenously produced in mammalian and is one of the most important regulators of vascular tone and blood pressure, concentration-response curves (CRC) for noradrenaline were performed to study the role of PGRN on the vascular tone control. Mesenteric arteries were incubated with rPGRN for 1h prior to noradrenaline curves in three different concentrations (100, 300, and 600ng/mL).
- *Protocol to study the endothelial factors*: CRC to noradrenaline was performed in the presence of cyclooxygenase 1 and 2 inhibitors (Indomethacin, 10µM), NOS inhibitor (L-NAME, 100µM), or the combination of indomethacin and L-NAME (to indirectly analyze the endothelium-derived hyperpolarizing factor, EDHF).
- *Protocol to study the receptors of PGRN*: Arteries were pre-incubated (30 minutes prior to rPGRN incubation) with EphrinA2 antagonist (ALW-II-4127, 5µM)^40^ or Sortilin1 inhibitor (AF38469, 40µM)^41^.
- *Protocol to study if deficiency in the PGRN affects vascular contractility*: CRC to noradrenaline was performed in mesenteric arteries from PGRN+/+ and PGRN-/- mice.

### Freshly endothelial cells isolation from mesenteric bed

Adapted from our previous publications^42, 43^. Male and female mice were sacrificed, and their mesenteric beds were excised and pooled, washed in PBS, and diced into small pieces, which were incubated in Dulbecco’s Modified Eagles Medium (DMEM; Gibco, Thermo Fisher Scientific, NH-USA) containing 10% fetal bovine albumin (FBS), 2mg/mL of collagenase II and 40 mg/ml dispase-II at 37°C for 1 hour while shaking. The cell suspension was vigorously vortexed and meshed through 40 µm nylon cell strainers (Fisherbrand, Thermo Fisher Scientific, NH-USA). After centrifugation, the cell pellet was resuspended in 1X PBS with 0.5% bovine serum albumin (BSA) and 2 mM ethylenediaminetetraacetic acid (EDTA). Endothelial cells were labeled with CD31-conjugated magnetic microbeads and sorted using magnetic separation LS columns (Miltenyi Biotech, German). RNA was isolated as described below and the purity of endothelial cells was checked by evaluating smooth muscle cells (αSMA) and endothelial cells (CD31 and eNOS) markers. Expression of EphrinA2 and Sortilin1 was evaluated in samples from male and female mice to analyze whether there might be any sex difference in PGRN receptors.

### Culture of endothelial cells

Mouse Mesenteric Endothelial Cells (mMEC) or human Mesenteric Endothelial Cells (hMEC) were purchased from Cell Biologics (Chicago, IL, USA) and were used to understand EphrinA2 and Sortlin1 are expressed in human and murine endothelial cells. Cells were maintained in Complete Mouse Endothelial Cell Medium (Cell Biologics, Chicago, IL, USA) containing Endothelial Cell Medium Supplement Kit (Cell Biologics, Chicago, IL, USA). Cells were used between passages 4-8.

### Protocol of endothelial cells treatment

Cells were treated with rPGRN (600ng/mL) at different times (0-60 minutes). To understand by which receptors PGRN induces endothelial cell activation. hMEC were treated with rPGRN with or without ALW-II-4127 (5µM) or AF38469 (40µM). Then, eNOS phosphorylation and NO levels were measured as described below. EphrinA2 and Sortilin1 protein expression was also analyzed in mMEC and hMEC.

### RT-PCR

mRNA from mesenteric bed and freshly isolated flow through and endothelial cells from mesenteric beds were extracted using RNeasy Mini Kit (Quiagen, Germantown, MD – USA). Complementary DNA (cDNA) was generated by reverse transcription polymerase chain reaction (RT-PCR) with SuperScript III (Thermo Fisher Waltham, MA USA). Reverse transcription was performed at 58 °C for 50 min; the enzyme was heat inactivated at 85 °C for 5 min, and real-time quantitative RT-PCR was performed with the PowerTrack™ SYBR Green Master Mix (Thermo Fisher, Waltham, MA USA). Sequences of genes as listed in supplementary table 1. Experiments were performed in a QuantStudio™ 5 Real-Time PCR System, 384-well (Thermo Fisher, Waltham, MA USA). Data were quantified by 2ΔΔ Ct and are presented by fold changes indicative of either upregulation or downregulation.

### Western Blot

mMEC and hMEC samples were directly homogenized using 2x Laemmli Sample Buffer and supplemented with 2-Mercaptoethanol (β-mercaptoethanol) (BioRad Hercules, California – USA). Proteins were separated by electrophoresis on a polyacrylamide gradient gel (BioRad Hercules, California – USA), and transferred to Immobilon-P poly (vinylidene fluoride) membranes. Non-specific binding sites were blocked with 5% skim milk or 1% bovine serum albumin (BSA) in tris-buffered saline solution with tween for 1h at 24 °C. Membranes were then incubated with specific antibodies overnight at 4 °C as described in supplementary table 2. After incubation with secondary antibodies, the enhanced chemiluminescence luminol reagent (SuperSignal™ West Femto Maximum Sensitivity Substrate, Thermo Fisher Waltham, MA, USA) was used for antibody detection.

### Nitric oxide measurement

Nitric oxide production was measured by a 4,5-Diaminofluorescein diacetate (DAF-2 DA) probe. Briefly, mMEC were treated with rPGRN (600ng/mL) for 60 minutes, with or without pre-incubation of ALW-II-4127 (5µM) or AF38469 (40µM), then cells were washed with PBS and stained with DAF-2 DA (5µM) for 30 minutes before analyze. Fluorescence intensity was analyzed in a fluorimeter (SpectraMax i3x Multi-Mode Microplate Reader) (Emission. 538 nm/Excitation. 485 nm). Bradykinin (10µM)^44^ was used as a positive control.

### Statistic

For comparisons of multiple groups, one-way or two-way analysis of variance (ANOVA), followed by the Tukey post-test was used. Differences between the two groups were determined using Student’s t-test. The vascular function data are expressed as a percentage of KCl (60mM)-induced maximal response (mN/KCL response). The concentration-response curves were fitted by nonlinear regression analysis. Maximal response (Emax) was determined. Analyses were performed using Prism 9.0 software (GraphPad). A difference was considered statistically significant when P ≤ 0.05.

## Results

### Hypertension is associated with high circulating levels of PGRN

Previous studies have shown that circulating PGRN is elevated in obesity^6, 7^, diabetes^8, 9^, and lipodystrophy^10–12^ – major causes of cardiovascular diseases. Here we observed that two different models of hypertension, Ang-II, or Aldo-treated mice, displayed increases in circulating PGRN (Fig.1A-D). However, why PGRN is enhanced in hypertension is fully unknown. In this study, we are investigating the importance of PGRN in maintaining blood pressure and vascular tone by using a global PGRN-deficient mouse and PGRN treatment.

**Figure 1.**
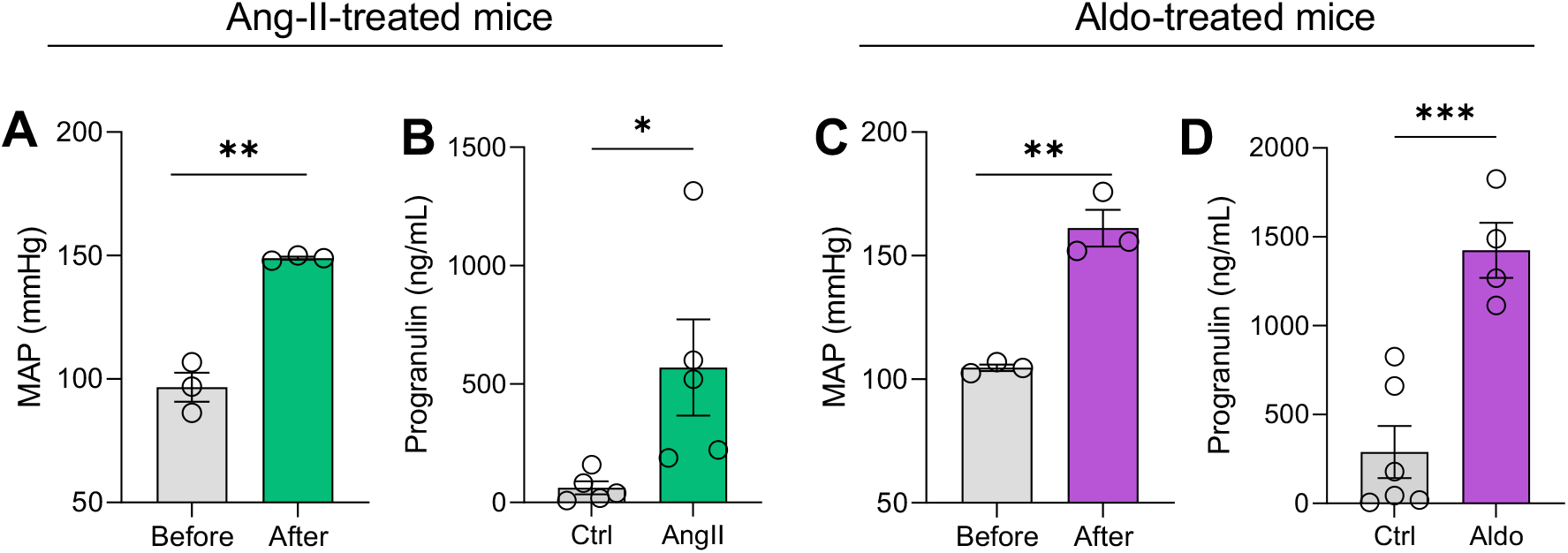
Hypertension is associated with high levels of circulating PGRN. Mean Arterial Pressure (MAP) and PGRN plasma levels in Ang-II treated mice (490ng/Kg/14 days) (A-B) or Aldo-treated mice (600ng/Kg/14 days) (C-D). Data are presented as Mean ± Standard Error of the Mean (SEM). N=3-6. *P<0.05 vs ctrl.

### PGRN deficiency does not affect vascular inflammation and structure but increases inflammatory circulating markers and sodium excretion

PGRN seems to have anti-inflammatory effects in different cells and organs and modulate cell growth^18, 20^, therefore we investigated whether lack of PGRN might affect the inflammatory profile, vascular fibrosis and hypertrophy, and sodium excretion. We observed that PGRN-/- mice do not present inflammation in mesenteric beds, but they display increases in circulating CCL2 (inflammatory chemokine), IL-13 (cytokine involved in allergic inflammation), and interferon-γ (inflammatory soluble cytokine), with no changes aortic remodeling (fibrosis, hypertrophy, and collagen amount) (Fig.2A-D). We also found that PGRN-/- mice excrete more sodium and a larger amount of urine compared to PGRN+/+ (Fig.2E), which are characteristics of hypertension^45–47^.

**Figure 2.**
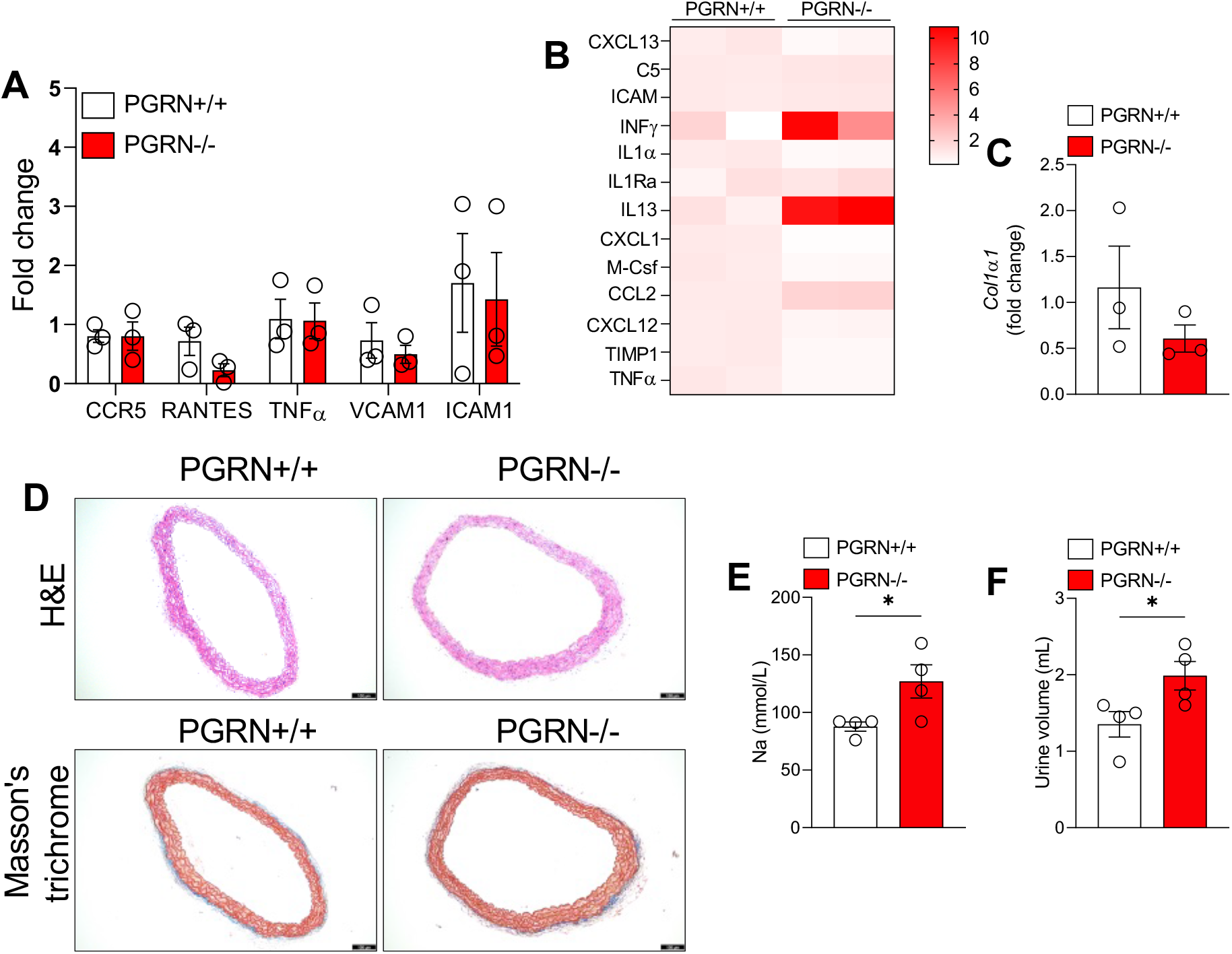
Deficiency in PGRN does not affect vascular inflammatory profile and aortic remodeling but increases circulating inflammatory markers and sodium and urine excretion. Inflammatory profile in mesenteric beds, measured by RT-PCR (A), and in plasma, measured by proteome profiler mouse cytokine array and presented as heat map (B), from PGRN+/+ and PGRN-/-. Collagen1a1 gene expression (C), H&E and Masson’s trichrome stains (D) in thoracic aortae from PGRN+/+ and PGRN-/-. 24 hours of urine sodium content (E) and volume (F) from PGRN+/+ and PGRN-/-. 11-13-weeks-old male mice were used. Scale bar= 100uM. Data are presented as Mean ± Standard Error of the Mean (SEM). N=3-4. *P<0.05 vs PGRN+/+. Cytokine profile was analyzed in a pool of 6 samples from each group and is represented as fold changes and represented in a het map.

### PGRN deficiency elevates blood pressure and increases vascular contractility

Via radiotelemetry, we found that lack of PGRN increases the MAP, SAP, and DBP with no effects on heart rate (Fig.3A-D). Changes in blood pressure do not seem to be dependent on sympathetic modulation because propranolol and mecamylamine affected similarly the blood pressure in PGRN+/+ and PGRN-/- (Fig.3E and F). Analysis of vascular contractility revealed that PGRN-/- mice did not show changes in KCl-induced contractility (Fig. 4A and D), but they presented higher vascular contractility compared to PGRN+/+ independent of sex since mesenteric arteries from male and female mice responded equally (Fig. 4B and D). Furthermore, an increase in contractility in males and females for noradrenaline (an adrenergic agonist) was not associated with changes in the expression of α-adrenergic receptors since no difference in α1a, α1b, and α1d was found (Fig. 4C and F). However, changes in sensitivity of adrenergic receptors have been previously reported in hypertension^48, 49^, therefore we cannot eliminate the chances of loss or gain of receptor activity including the α and β-adrenergic receptors family.

**Figure 3.**
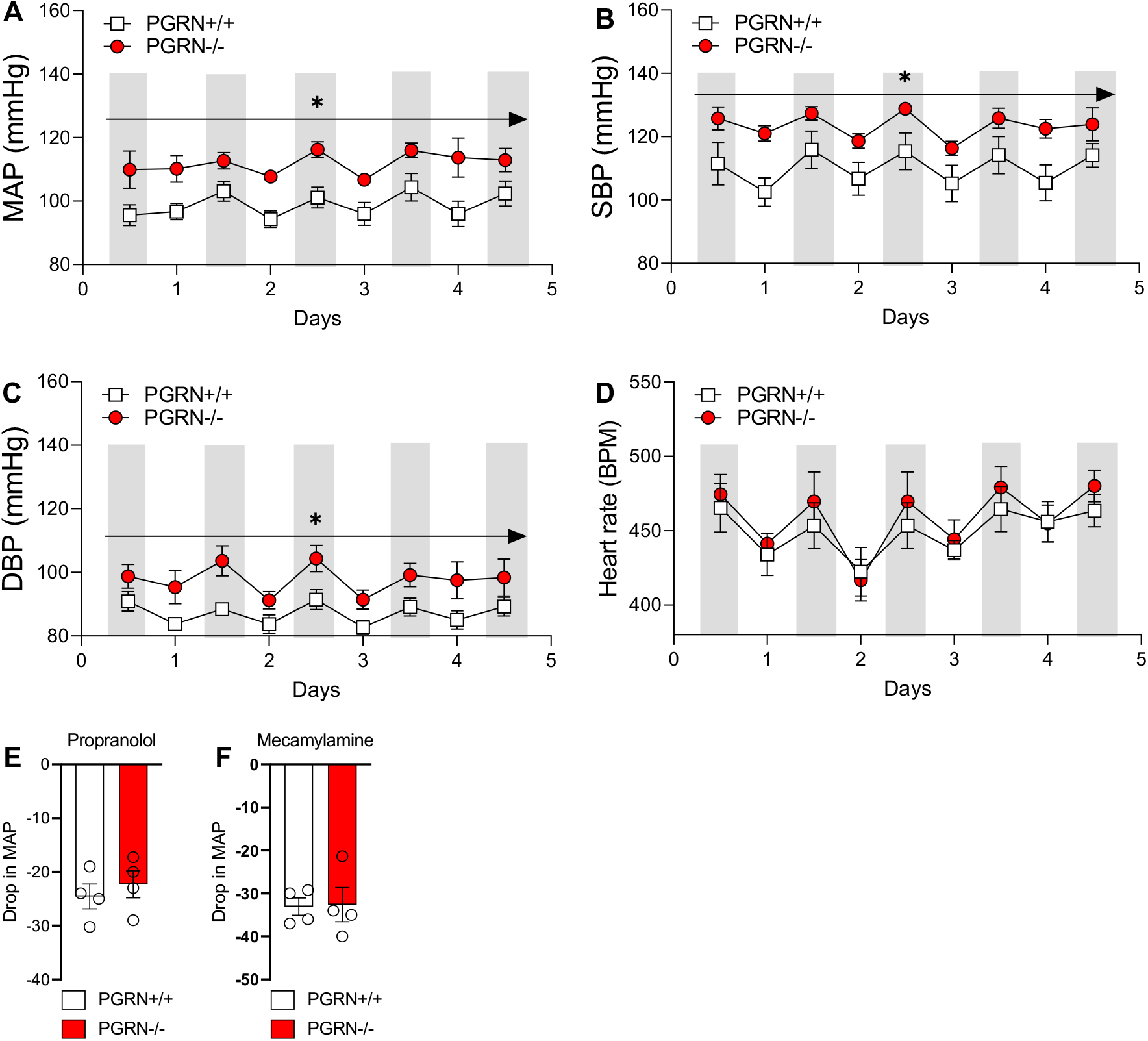
Deficiency in PGRN triggers high blood pressure. Mean arterial pressure (MAP), systolic blood pressure (SBP), diastolic blood pressure (DBP), and heart rate measured via radiotelemetry in male (11-13-weeks-old) PGRN+/+ and PGRN-/- (A-D). Effects of propranolol, 6 mg/kg (E) and mecamylamine, 5 mg/kg (F) on MAP. Gray bars represent nighttime in telemetry. Data are presented as Mean ± Standard Error of the Mean (SEM). N=3-4. *P<0.05 vs PGRN+/+.

**Figure 4.**
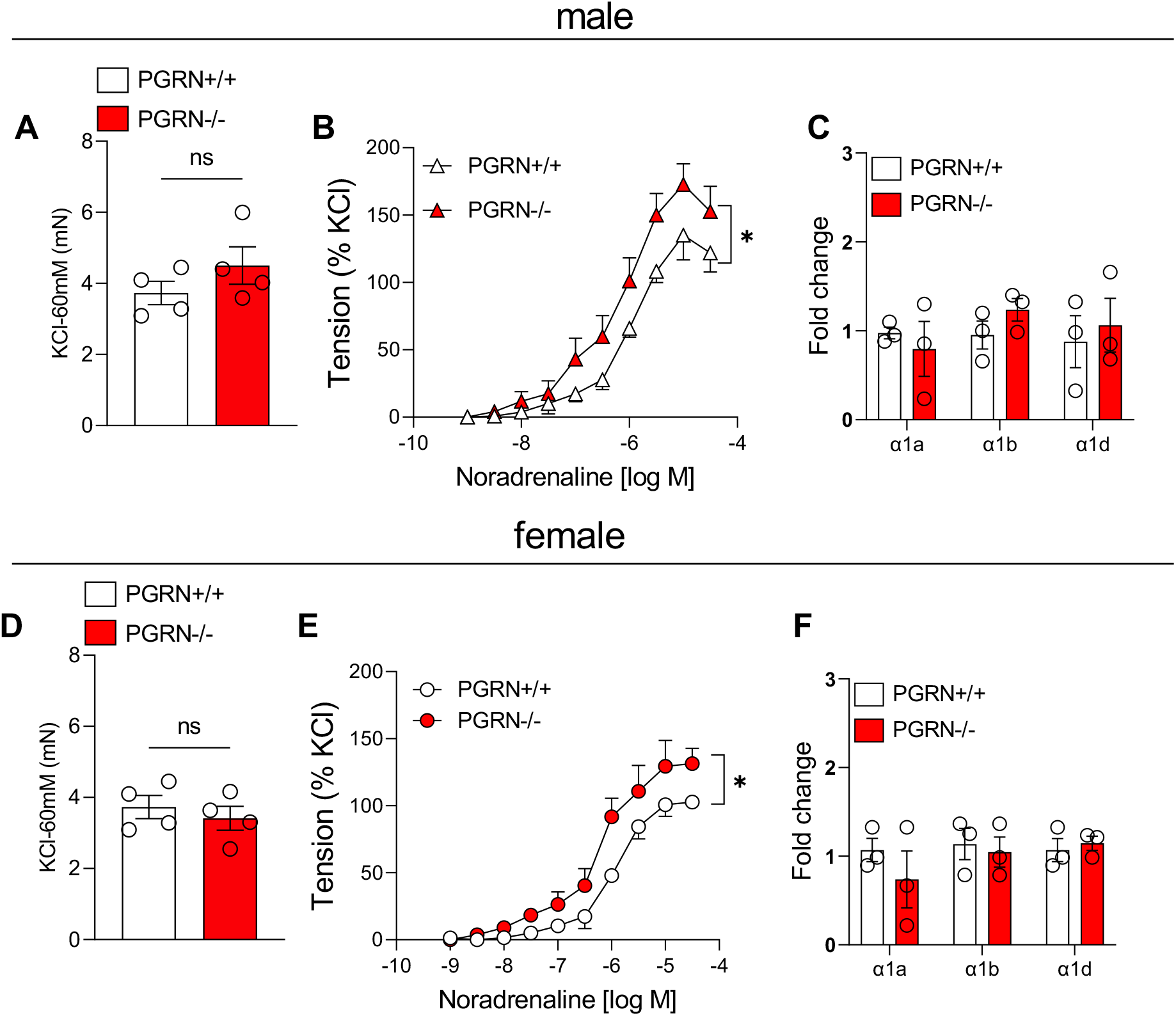
Deficiency in PGRN triggers vascular dysfunction. KCl (60mM) response in mesenteric arteries (2^nd^ order) from male (A) and female (C) PGRN+/+ and PGRN-/- mice. Concentration responses curves (CRC) to noradrenaline in mesenteric arteries from male (B) and female (E) PGRN+/+ and PGRN-/- mice. α1-adrenergic gene expression in mesenteric beds from male (C) and female (F) PGRN+/+ and PGRN-/- mice. Data are presented as Mean ± Standard Error of the Mean (SEM). N=3-4. *P<0.05 vs PGRN+/+.

### PGRN replacement restores blood pressure and vascular contractility

To understand whether circulating PGRN maintains blood pressure and vascular function we sought to restore PGRN levels by treating PGRN-/- mice with rPGRN (20ug/day/7 days, via osmotic mini-pump)^13^. Treatment with rPGRN increased the PGRN levels in PGRN-/-, but not restored as PGRN+/+ mice, furthermore in PGRN+/+ mice treatment with rPGRN slightly elevated PGRN levels (Fig.5A). In addition, rPGRN treatment decreased the vascular contractility in male and female PGRN-/- mice (Fig.5B and 4C), as well as recovered the MAP, SBP, and DBP, with no significant effects in PGRN+/+. No effect was observed in heart rate (Fig.5D-G). These data suggest that circulating PGRN helps maintain physiological blood pressure levels and that arteries and blood pressure from PGRN-/- mice are more sensitive to rPGRN treatment.

**Figure 5.**
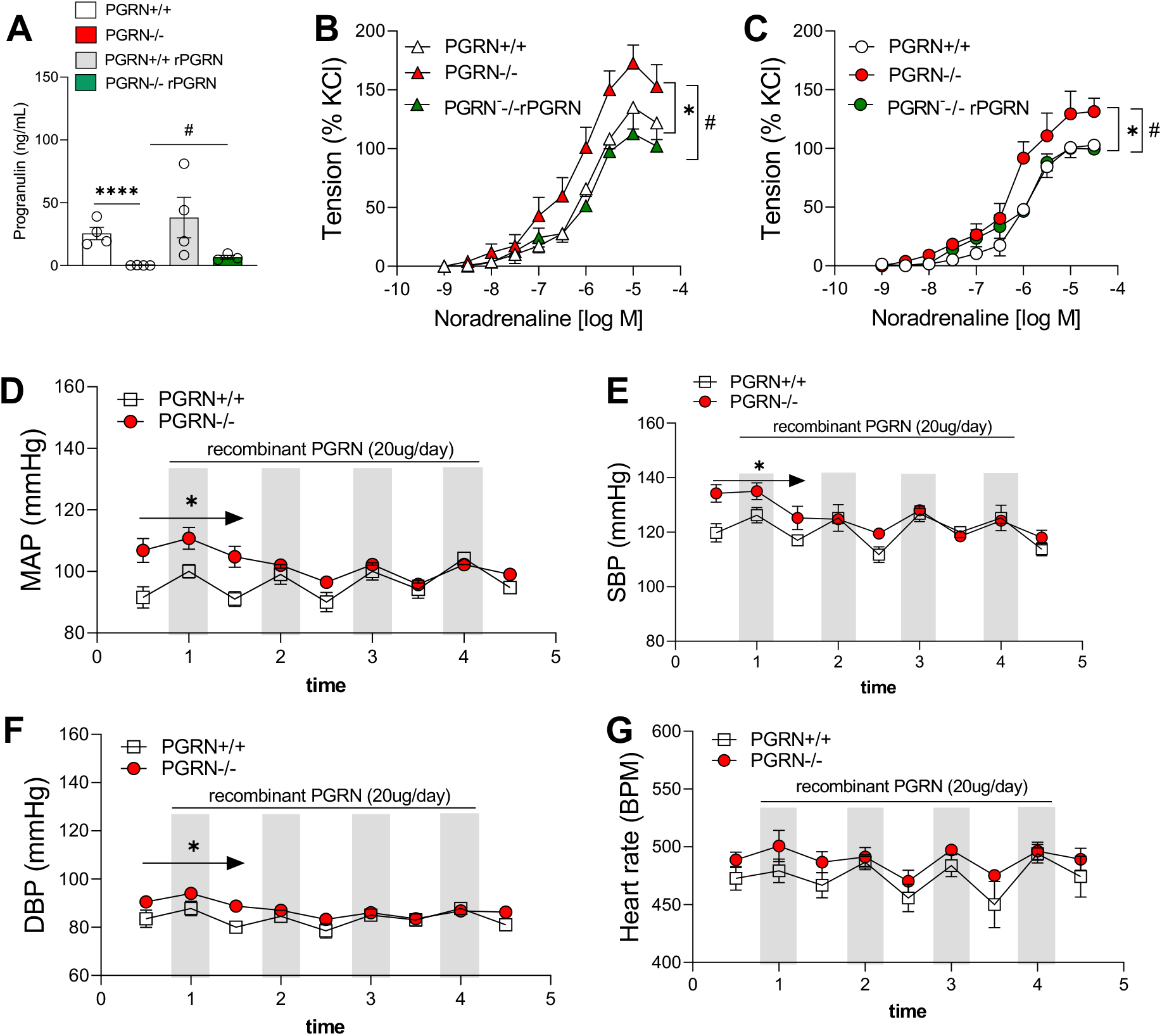
PGRN treatment restores vascular function and blood pressure in PGRN deficient mice. Circulating PGRN levels (A) and concentration responses curves (CRC) to noradrenaline in mesenteric arteries from male (B) and female (C) PGRN+/+ and PGRN-/- mice treated or not with rPGRN (20ug/day/7-days). Mean arterial pressure (MAP) (D), systolic blood pressure (SBP) (E), diastolic blood pressure (DBP) (F), and heart rate (G) measured via radiotelemetry in male (11-13-weeks-old) PGRN+/+ and PGRN-/- (A-D) treated with rPGRN (20ug/day/7-days). Gray bars represent nighttime in telemetry. Data are presented as Mean ± Standard Error of the Mean (SEM). N=3-4. *P<0.05 vs PGRN+/+ and ^#^P<0.05 vs PGRN-/-.

### PGRN attenuates vascular contractility in males and females dependent on EphrinA2 and Sortilin1

To study the mechanisms whereby PGRN exerts its anti-contractility effects, we treated mesenteric arteries from male and female control mice (PGRN+/+) for 1h prior to noradrenaline CRC. We found that 600ng/ml of PGRN, but not 100 or 300ng/mL, decreased vascular contractility in male and female mice (Fig.6A and B).

**Figure 6.**
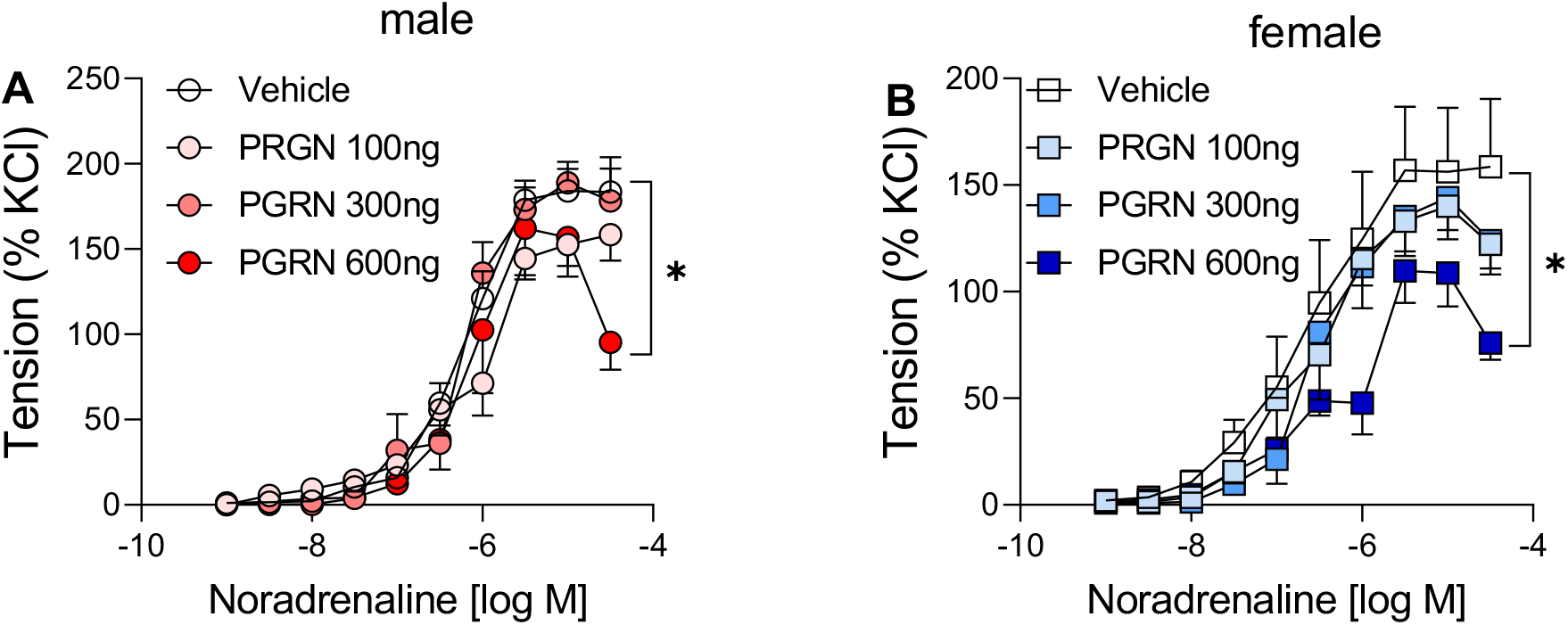
PGRN incubation attenuates vascular contractility. Effects of PGRN (100, 300, and 600ng/mL/1h) on concentration responses curves (CRC) to noradrenaline in mesenteric arteries (2^nd^ order) from male (A) and female (B) C57BL6/J mice (11-13- weeks-old). Data are presented as Mean ± Standard Error of the Mean (SEM). N=4-6. *P<0.05 vs vehicle.

Next, we evaluated by which receptor PGRN exerts its anti-contractile effect via pharmacologically blocking two known receptors for PGRN: EphrinA2 and Sortilin1. Antagonism of EphrinA2, with ALW-II-4127 (5uM), or Sortilin1, with AF38469, (40uM) prevented the rPGRN effects in mesenteric arteries from male and female mice (Fig. 7A-D).

**Figure 7.**
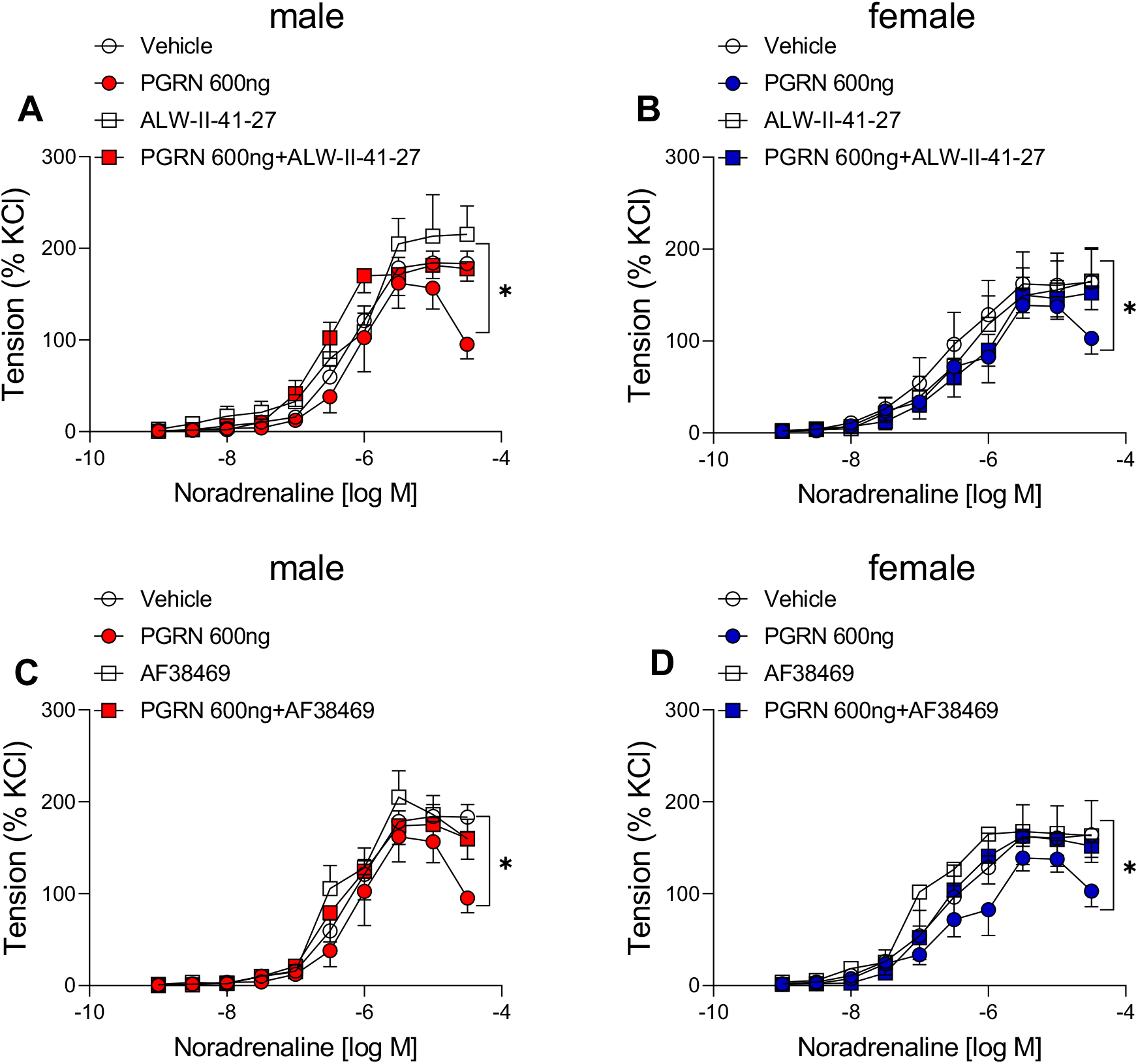
PGRN incubation attenuates vascular contractility dependent on EphrinA2 and Sortilin1. Concentration responses curves (CRC) to noradrenaline in mesenteric arteries (2^nd^ order) from male (A and C) and female (B and D) C57BL6/J mice (11-13- weeks-old) in presence of PGRN (600ng/mL/1h) with or without EphrinA2 antagonist (ALW-II-4127, 5µM) (A and B) or Sortilin1 inhibitor (AF38469, 40µM) (C and D). Data are presented as Mean ± Standard Error of the Mean (SEM). N=4-6. *P<0.05 vs arteries treated with rPGRN.

### PGRN reduces vascular contractility via nitric oxide production

To study the molecular mechanisms whereby PGRN reduces adrenergic contractility in male and female mice, we used different pharmacological tools to block endothelial-derived factors formation including nitric oxide, prostaglandins production, and EDHF. We found that L-NAME (NOS inhibitor) blunted the difference in noradrenaline response caused by PGRN incubation in mesenteric arteries from male and female mice (Fig. 8A and D), whereas indomethacin (COXs inhibitor) only affected the response in arteries incubated with the vehicle but did not impact the PGRN effects in both sexes (Fig. 8B and E). Furthermore, L-NAME and indomethacin combination did not change any response in mesenteric arteries from male and female mice (Fig. 8C and 8E). These findings suggest that PGRN exerts its anti-contractile effects via modulating nitric oxide formation and/or attenuating contractile prostaglandins.

**Figure 8.**
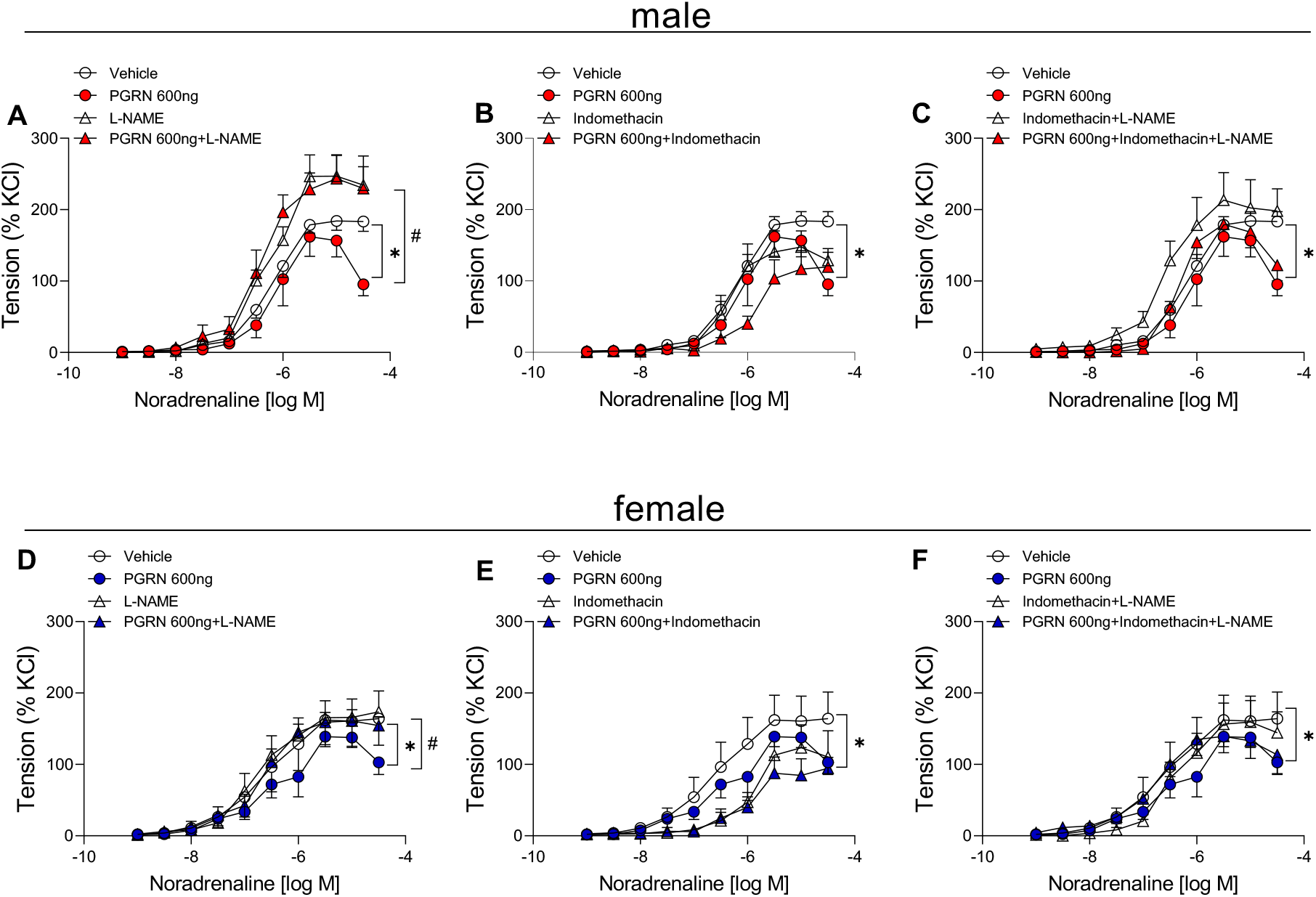
PGRN incubation attenuates vascular contractility dependent on nitric oxide. Concentration responses curves (CRC) to noradrenaline in mesenteric arteries (2^nd^ order) from male (A-C) and female (D-F) C57BL6/J mice (11-13-weeks-old) in presence of PGRN (600ng/mL/1h) with or without NOS inhibitor (L-NAME, 100µM) (A and D), cyclooxygenase inhibitor (indomethacin, 10µM) (B and D), or L-NAME+ indomethacin (C and F). Data are presented as Mean ± Standard Error of the Mean (SEM). N=4-6. *P<0.05 vs arteries treated with rPGRN; ^#^P<0.05 vs L-NAME.

Finally, we established a protocol of fresh isolation of endothelial cells from mesenteric beds by using CD31^+^ microbeads (Fig. 9A) to measure the expression of EphrinA2 and Sortilin1 expression in endothelial cells from male and female mice to analyze any sex difference in PGRN receptors expression. We first characterized the purity of endothelial cells by measuring the gene expression of αSMA (a marker of the smooth muscle cell) and CD31 and eNOS (markers of endothelial cells) in flow through and endothelial cells, we found a striking reduction in αSMA along with a clear increase in CD31 and eNOS in endothelial cells versus flow through (Fig. 9B). Analyze of the expression of EphrinA2 and Sortilin1 in freshly isolated endothelial cells from male and female revealed no difference between the sexes (Fig. 9C).

**Figure 9.**
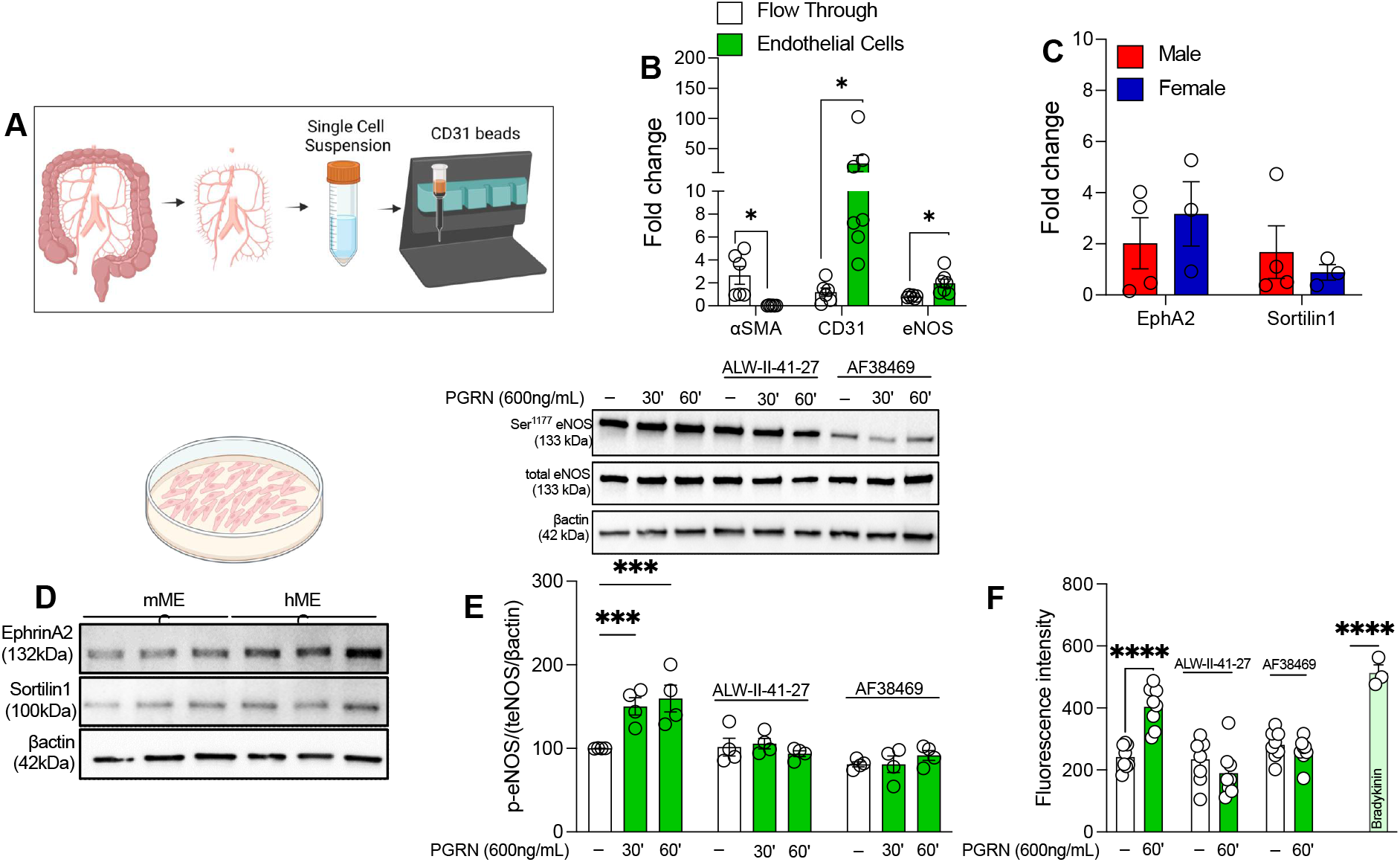
PGRN activates eNOS in mesenteric endothelial cells via EphrinA2 and Sortilin1. Schematic of endothelial cells isolation from mesenteric bed (A). Smooth muscle cell (aSMA) and endothelial cell markers (CD31 and eNOS) in freshly isolated endothelial cells and flow through from mesenteric bed measured by RT-PCR (B). EphrinA2 and Sortilin1 gene expression in freshly isolated endothelial cells from male and female mice (C). Expression of EphrinA2 and Sortilin1 in mouse mesenteric endothelial cells (mMEC) and human mesenteric endothelial cells (hMEC) (D). Effects of rPGRN (600ng/mL for 30 and 60 minutes) on eNOS phosphorylation (Ser^1177^) in mMEC (E). Effects of rPGRN on nitric oxide formation measured by DAF-AM (F). Bradykinin (10µM) was used as a positive control. Experiments were performed in presence or absence of EphrinA2 antagonist (ALW-II-4127, 5µM) or Sortilin1 inhibitor (AF38469, 40µM). Data are presented as Mean ± Standard Error of the Mean (SEM). N=3-8. *P<0.05 vs flow through or *P<0.05 vs vehicle in experiments of mMEC treated with rPGRN.

In this study, we particularly focused on understanding whether PGRN induces nitric oxide formation in mesenteric endothelial cells. Thus, we first analyzed if mesenteric endothelial cells from mice and humans express EphrinA2 and Sortilin1, as demonstrated in figure 9D, both receptors are well-expressed in mouse and human cells. Next, we analyzed whether rPGRN induces eNOS activation and nitric oxide formation in mMEC. PGRN triggered eNOS activation (phosphorylation of Ser^1177^ residue) and elevated nitric oxide production, which were prevented by blocking EphrinA2 and Sortilin1 (Fig. 9E and F). Bradykinin was used as a positive control of nitric oxide production. Therefore, we can hypothesize that PGRN modifies the vascular tone and blood pressure by controlling the synthesis of nitric oxide and prostanoids, which seem to be dependent on EphrinA2 and Sortilin1.

## Discussion

Although PGRN is extensively researched in neurodegenerative disorders^18, 19, 50^, little is known about its significance in vascular biology and blood pressure regulation. Our study is the first to extensively examine how circulating PGRN affects blood pressure regulation and vascular tone. As a result, we were able to show that, under physiological conditions, circulating PGRN aids in regulating vascular resistance and possibly blood pressure maintenance. Moreover, high levels of PGRN associated with HBP may act as a defensive mechanism to help lower blood pressure.

Lastly, it appears that these protective processes depend on EphrinA2, Sortlin1, and nitric oxide. According to these findings, individuals with low PGRN levels may be more susceptible to cardiovascular events, whereas chronic PGRN replacement treatment is not only safe but also advantageous for the cardiovascular system in individuals with a deficiency in PGRN. Moreover, the therapy of HBP with PGRN may be an interesting therapeutic route because it likely modulates nitric oxide formation.

Although PGRN effects have been studied in other diseases, little is known about how it may affect cardiovascular physiology or pathophysiology. For instance, PGRN-deficient mice exhibit worse cardiorenal phenotype in a hyperhomocysteinemia diet^51^, greater renal injury in the diabetes model^15^, and accelerated heart hypertrophy on an age-dependent basis^24^. More recently, Gerrits et al ^50^ revealed that the neurovascular unit is severely affected in PGRN-associated frontotemporal dementia. Although these studies implicate that lack of PGRN induces cardiovascular and renal dysfunction or predisposes a worse phenotype in different conditions, the role of PGRN deficiency in controlling blood pressure is not well-explored. Previous findings reported that PGRN is elevated in plasma from hypertensive patients^52, 53^, but why and which cells are producing this PGRN are fully unclear. In our two models of hypertension, circulating PGRN was strikingly high. However, we found that aldosterone plus saline treatment produced a further increase in PGRN compared to Ang-II-treatment indicating that there might be a renal contribution to such discrepancy, which could be mediated by volume-dependent hypertension mechanisms since aldosterone and salt is a model of increased blood volume. Further studies are necessary to evaluate this inconsistency between the hypertension models, as well as to understand which cells are mostly producing PGRN in hypertension.

We also described here that PGRN maintains the blood pressure and vascular tone by observing that global PGRN deficient mice display elevated blood pressure and increased vascular contractility followed by a large amount of sodium excretion and circulating inflammatory profile. Our data differ from previous findings, where the authors found, via echocardiography^24, 51^, that deficiency in PGRN does not affect blood pressure. Possibly our blood pressure recorded via telemetry was more sensitive to detect any small changes in blood pressure. Interestingly, we could restore the blood pressure in PGRN-/- mice by returning circulating PGRN for 7 consecutive days. Surprisingly, rPGRN treatment (20ug/day) did not achieve PGRN levels as the control group but only increased it. We postulate that PGRN might be degraded, cleaved by proteases, or internalized by different cells^18–20, 54^, thus reducing the circulating availability. Further investigation is needed to dissect such an event.

A study published by Dr. Yamawaki’s group in 2017^55^ revealed that isolated superior rat mesenteric artery rings incubated with PGRN (10–100 ng/mL) show an increased sensitivity to acetylcholine suggesting that PGRN is physiologically relevant to keep the vascular tone. Aligned with these findings, we observed that deficiency in PGRN triggers vascular dysfunction, characterized by the elevated response to noradrenaline, which is blunted by supplementing PGRN-/- mice with rPGRN. Furthermore, we found that 600ng/mL (6 times more PGRN than shown previously^55^), attenuates noradrenaline response in mesenteric arteries from male and female mice. Therefore, we can suggest that circulating PGRN maintains vascular tone and blood pressure and, in hypertension models, circulating PGRN is likely elevated as a compensatory mechanism to promote a decrease in vascular resistance and subsequently decrease the blood pressure.

Several PGRN receptors have been identified, including EphrhinA2^56^ and Sortilin1^19^. EphrhinA2 belongs to a family of receptor tyrosine kinases which is crucial for migration, and vascular and epithelial development^56, 57^. Studies have shown that PGRN can bind to EphrinA2 and induce AKT activation and angiogenesis^56^. While Sortlin1 is a sorting receptor that can reduce PGRN bioavailability by capturing and transporting it to the lysosome for destruction^58^, not all PGRN proceeds along the lysosome pathway; instead, some may move to an unknown additional cellular compartment^58^. Here, we found that blocking EphrinA2 or Sortilin1 blunts the anti-contractile action brought on by PGRN. These findings indicate that a possible crosstalk between EphrinA2 and Sortilin1 may exist and that further research is necessary to unravel such communication. Perhaps Sortlin1 might regulate some intracellular pathways, which can interfere with EphrinA2 receptor activation, e,g. AKT pathway^56^ or even a direct modulation of intracellular protein–tyrosine kinase domain by Sortilin1 pathway. Since we found that PGRN affects vascular contractility in males and females, we investigated whether EphrinA2 or Sortilin1 expression is similar in male and female mesenteric endothelial cells, we discovered that both male and female mesenteric endothelial cells express these two PGRN receptors equally.

Endothelial cells are responsive to PGRN effects^33, 56^. PGRN can directly protect the vascular endothelium against atherosclerotic environment by eNOS and NFkB^33^, induce capillary morphogenesis and GRN autoregulation, and influences the growth and development of blood vessels^54, 56, 59^. We found that the anti-contractile effect of PGRN is mediated by nitric oxide formation since blocking NOS blunted the difference between arteries incubated or not with rPGRN. Moreover, such an effect seems to be sex independent because arteries from male and female mice responded similarly. Also, we discovered that PGRN may alter vasoconstrictor prostanoids. This conclusion may be drawn from the observation that only naive arteries, not PGRN-treated arteries, responded to cyclooxygenase inhibitors. To better understand the methods through which PGRN modifies the cyclooxygenase pathway, more research is required. Since double inhibition (NOS and cyclooxygenase) did not affect on any PGRN response, our data further imply that EDHF does not appear to be involved in the effects of PGRN.

As mentioned previously, PGRN can modulate nitric oxide production in the endothelium^33^. To confirm whether PGRN induces nitric formation via EphrinA2 or Sortilin1 we took advantage of isolated mesenteric endothelial cells. Firstly, we demonstrated that mesenteric endothelial cells from mice and humans express both receptors, then we observed that PGRN leads to eNOS activation and nitric oxide formation via EphrinA2 or Sortilin1 activation. Thus, we can suggest that PGRN modulates the vascular tone via EphrinA2 or Sortilin1 and eNOS activation.

In conclusion, this work offers the first proof that PGRN aids in controlling blood pressure and vascular tone through the production of nitric oxide and EphrinA2 or Sortilin1. Moreover, we have shown that PGRN exerts an anti-contractile effect in both male and female mice via similar mechanisms. Together, these findings suggest that PGRN deficiency may contribute to vascular dysfunction, and hypertension, or predispose patients to a more severe cardiovascular outcome. Therefore, PGRN may provide a novel treatment strategy for lowering blood pressure and restoring vascular tone. However, further research is required to comprehend 1. the relationship between EphrinA2 and Sortilin1, 2. the effects of PGRN supplementation in established high blood pressure, and 3. the toxicological effects of PGRN specially on hepatic integrity.

**Acknowledgments, Sources of Funding, & Disclosures**

## Abbreviations

Progranulin: PGRN
Nitric oxide synthase: NOS
Endothelial NOS: eNOS
High blood pressure: HBP
Endothelium-derived relaxing factors: EDRF
Endothelium-derived hyperpolarizing factor: EDHF
Endothelium-derived contracting factors: EDCF
Protein kinase B: AKT
Nuclear factor-κB: NFkB
Angiotensin II: Ang-II
Aldosterone: Aldo
Phosphate-buffered saline: PBS
Paraformaldehyde: PFA
Mouse Mesenteric Endothelial Cells: mmec
Human Mesenteric Endothelial Cells: hmec
2-Mercaptoethanol: β-mercaptoethanol
Radioimmunoprecipitation assay buffer: RIPA
α Smooth Muscle Actin: αSMA
Platelet endothelial cell adhesion molecule: CD31

## Acknowledgments

This work was supported by: NHLBI-R00 (R00HL14013903), AHA-CDA (CDA857268), Vascular Medicine Institute, the Hemophilia Center of Western Pennsylvania Vitalant, in part by Children’s Hospital of Pittsburgh of the UPMC Health System, and startup funds from University of Pittsburgh to TBN. Schematic in figure 9 and graphical abstract were designed using BioRender.

