## Supplemental tables for "Progranulin maintains blood pressure and vascular tone dependent on EphrinA2 and Sortilin1 receptors and eNOS activation"

**Table 1. List of primers**

| Primer | Sequence | |
|---|---|---|
| $\alpha$SMA | FW | TGCTGACAGAGGCACCACTGAA |
| | RV | CAGTTGTACGTCCAGAGGCATAG |
| CD31 | FW | CCAAAGCCAGTAGCATCATGGTC |
| | RV | GGATGGTGAAGTTGGCTACAGG |
| eNOS | FW | CGCAAGAGGAAGGAGTCTAGCA |
| | RV | TCGAGCAAAGGCACAGAAGTGG |
| EphrinA2 | FW | CAAAGTGCACGAGTTCCA |
| | RV | CTCCTGCCAGTACCAGAAGC |
| Sortilin1 | FW | CTACTCCATCCTGGCAGCCAAT |
| | RV | CTACTCCATCCTGGCAGCCAAT |
| $\alpha$-adrenergic 1a | FW | GGCTGGAGCATGGGTATATG |
| | RV | CTGCCATTCTTCCTCGTGAT |
| $\alpha$-adrenergic 1b | FW | GGCATTGTAGTCGGAATGTTCATC |
| | RV | CTGTTGAAGTAGCCCAGCCAGA |
| $\alpha$-adrenergic 1d | FW | GCTGGTTCCCCTTTTTCTTC |
| | RV | ATTGAAGTAGCCCAGCCAGA |
| GAPDH | FW | GAGAGGCCCTATCCCAACTC |
| | RV | TCAAGAGAGTAGGGAGGGCT |

Primers were purchased from Integrated DNA Technologies

**Table 2. List of antibodies**

| Antibody | Catalog number | Company | Concentration |
|---|---|---|---|
| Sortilin1 | 20681 | Cell Signaling | 1:1000 |
| EphrinA2 | sc-398832 | Santa Cruz | 1:500 |
| Ser1177 eNOS | 612393 | BD Biosciences | 1:1000 |
| eNOS | 610296 | BD Biosciences | 1:1000 |
| βactin | A3854 | Sigma | 1:20000 |
